# Quantification of Structural Brain Connectivity via a Conductance Model

**DOI:** 10.1101/415489

**Authors:** Aina Frau-Pascual, Morgan Fogarty, Bruce Fischl, Anastasia Yendiki, Iman Aganj, ^†^for the Alzheimer’s Disease Neuroimaging Initiativey

**Author notes:** Corresponding author, Email address (Aina Frau-Pascual). Data used in preparation of this article were obtained from the Alzheimer’s Disease Neuroimaging Initiative (ADNI) database (adni.loni.usc.edu). As such, the investigators within the ADNI contributed to the design and implementation of ADNI and/or provided data but did not participate in analysis or writing of this report. A complete listing of ADNI investigators can be found at: http://adni.loni.usc.edu/wp-content/uploads/how to apply/ADNI Acknowledgement List.pdf.

## Abstract

Connectomics has proved promising in quantifying and understanding the effects of development, aging and an array of diseases on the brain. In this work, we propose a new structural connectivity measure from diffusion MRI that allows us to incorporate direct brain connections, as well as indirect ones that would not be otherwise accounted for by standard techniques and that may be key for the better understanding of function from structure. From our experiments on the Human Connectome Project dataset, we find that our measure of structural connectivity better correlates with functional connectivity than streamline tractography does, meaning that it provides new structural information related to function. Through additional experiments on the ADNI-2 dataset, we demonstrate the ability of this new measure to better discriminate different stages of Alzheimer’s disease. Our findings suggest that this measure is useful in the study of the normal brain structure, and for quantifying the effects of disease on the brain structure.

## 1. Introduction

Neurological diseases affect a large and increasing portion of the population. Recent studies on brain structural and functional connectivity have focused on the impact of various diseases on the brain connections. Different neurological diseases have been shown to alter brain connectivity: Alzheimer’s disease (AD) (Rose et al., 2000; Buckner et al., 2005, 2009; Prasad et al., 2015; Yu et al., 2017), late-life major depressive disorder (Smagula and Aizenstein, 2016), epilepsy (Taylor et al., 2015), Parkinsons’s disease (Canu et al., 2015; Shah et al., 2017), schizophrenia (Arbabshirani et al., 2013; Cabral et al., 2013) and other psychosis (van Dellen et al., 2015; Mighdoll et al., 2015). Brain connectivity analysis has also proven to be useful in the study of the effects of aging on the brain (Damoiseaux, 2017; Wu et al., 2013; Salat, 2011; Fjell et al., 2016).

Brain structural and functional connectivities, as measured by diffusion-weighted MRI (dMRI) and resting-state functional MRI (rs-fMRI), respectively, reveal distinct features (Park and Friston, 2013). While structural connectivity reflects the white-matter axon bundles, functional connectivity measures the temporal correlation of blood oxygenation changes. Functional connectivity is based on the premise that brain regions that are activated synchronously, and therefore undergo similar oxygenation and deoxygenation variations, are related. However, functional connectivity may depend on the subject state (e.g., time of day, alertness, caffeine levels). Structural connectivity, on the other hand, is independent of the subject state, and can even be acquired *ex vivo*. Despite the fundamental differences in the processes that we observe via these two distinct measurements, structural and functional connectivities have been shown to be correlated (Damoiseaux and Greicius, 2009; Huang and Ding, 2016). Strong functional connections are, however, commonly observed between regions with no direct structural connection (Koch et al., 2002; Honey et al., 2009). Part of this variance has been shown to be due to the impact of indirect structural connections (Honey et al., 2009; Van Den Heuvel et al., 2009; Deligianni et al., 2011), i.e. between regions that are physically connected through multiple direct fiber bundles. Such indirect structural connections are usually considered only during the network analysis (Sporns, 2013), but not at the fiber tracking (tractography) step.

Differences in brain connectivity patterns between healthy and diseased populations are an indicator of change in the brain “wiring” and function due to the disease. In particular, AD has been found to impact connectivity (Rose et al., 2000; Buckner et al., 2005, 2009; Prasad et al., 2015; Yu et al., 2017). The progressive neurodegeneration suffered in AD, possibly caused by the spread and accumulation of misfolded proteins along structural connections in the brain (Iturria-Medina et al., 2014), affects the functional networks detected with rs-fMRI (Buckner et al., 2005, 2009; Yu et al., 2017) and the brain connections reconstructed with dMRI (Rose et al., 2000; Prasad et al., 2015). Accurately modeling structural connectivity could therefore reveal the effects of AD progression in white-matter degeneration.

The purpose of the present work is to derive a new structural-connectivity measure that considers all possible pathways, direct and indirect, with the aim of accessing more of the information that is often only available through functional connectivity. To that end, we propose a connectivity computation method that, by exploiting the well-studied mathematics of electrical circuits, models the white-matter pathways imaged via dMRI. Although macroscopic electrical models have been exploited in the context of brain connectivity (Chung et al., 2012; Aganj et al., 2014; Chung et al., 2017), we take a different approach and use partial differential equations (PDEs). Our method accounts for all possible white-matter pathways and is solved globally. It also does not require any parameter tuning and does not suffer from local minima, thereby producing consistent results with respect to the input data.

We evaluate the performance of the proposed conductance measure to test two different hypotheses, using a different dataset for each. First, we check the relationship between structural and functional connectivities, to test if our structural connectivity measure, which includes indirect connections, is more related to functional connectivity than standard measures are. Our results on 100 subjects of the WashU-UMN Human Connectome Project (HCP) (Van Essen et al., 2013) dataset support this hypothesis. Second, we test if our measure can classify healthy and diseased populations. Through experiments on data from the second phase of Alzheimer’s Disease Neuroimaging Initiative (ADNI-2) (Jack et al., 2008; Beckett et al., 2015), our connectivity measure proves useful in finding differences between the different stages of AD. This could lead to the discovery of new imaging biomarkers that help us better identify the disease stage, predict the onset of disease from connectivity patterns, and deepen our insight into the human brain and how it is affected by disease.

We present our conductance method for inferring a new structural brain connectivity measure from dMRI in Section 2 and put it into perspective with respect to standard tractography methods. We describe our analysis pipeline for the two datasets in Section 3 and present our results in Section 4. A discussion and conclusions follow in Sections 5 and 6, respectively. A preliminary abstract of this work has previously been presented (Frau-Pascual et al., 2018).

## 2. Methods

Brain connectivity has previously been modeled with the help of the established mathematical framework for macroscopic electrical circuits (Chung et al., 2012; Aganj et al., 2014; Chung et al., 2017). In this work, we make use of a combination of *differential* circuit laws, resulting in an equation similar to the heat equation proposed by O’Donnell et al. (2002). As illustrated in Fig. 1, we assign to each image voxel a local anisotropic conductivity value *D*, which is the diffusion tensor computed from dMRI (Basser et al., 1994; Tuch et al., 2001). (See Appendix B for a formulation to use higher-order diffusion models.) By solving the partial differential equation (PDE) (Haus and Melcher, 1989),

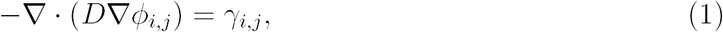

for a certain current configuration *γ*_*i,j*_ between a pair of source (*i*) and sink (*j*) voxels (see below), we find the potential map *ϕ*_*i,j*_ for that specific configuration. ∇ and∇ are the gradient and the divergence operators, respectively. The above PDE is the result of combining 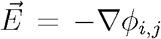 (due to Faraday’s law), 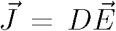 (Ohm’s law), and 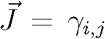 (Kirchhoff’s differential current law or continuity equation), where 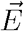is the electric field and 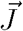 is the current density. The potential map *ϕ*_*i,j*_, an example of which is shown in Fig. 1(b), describes how a current would diffuse from a source point to a sink point following the orientational information provided by the gradient of the map and the diffusion tensors. It is nonetheless important to note that these potentials cannot be directly interpreted as connectivity. Therefore, in contrast to O’Donnell et al. (2002), we further compute the electric conductance between each pair of voxels from potential maps, to which all diffusion paths between the pair contribute (see Fig. 1(c)). The conductance, computed as the ratio of the total passed current to the potential difference, reflects the ease with which the current diffuses from the source to the sink, which we here use it as a measure of connectivity (more details provided below). The same measure can also be computed between a pair of regions of interest (ROIs) instead of voxels, by distributing the currents *γ* among the ROI voxels (see Fig. 1(c)).

**Figure 1:**
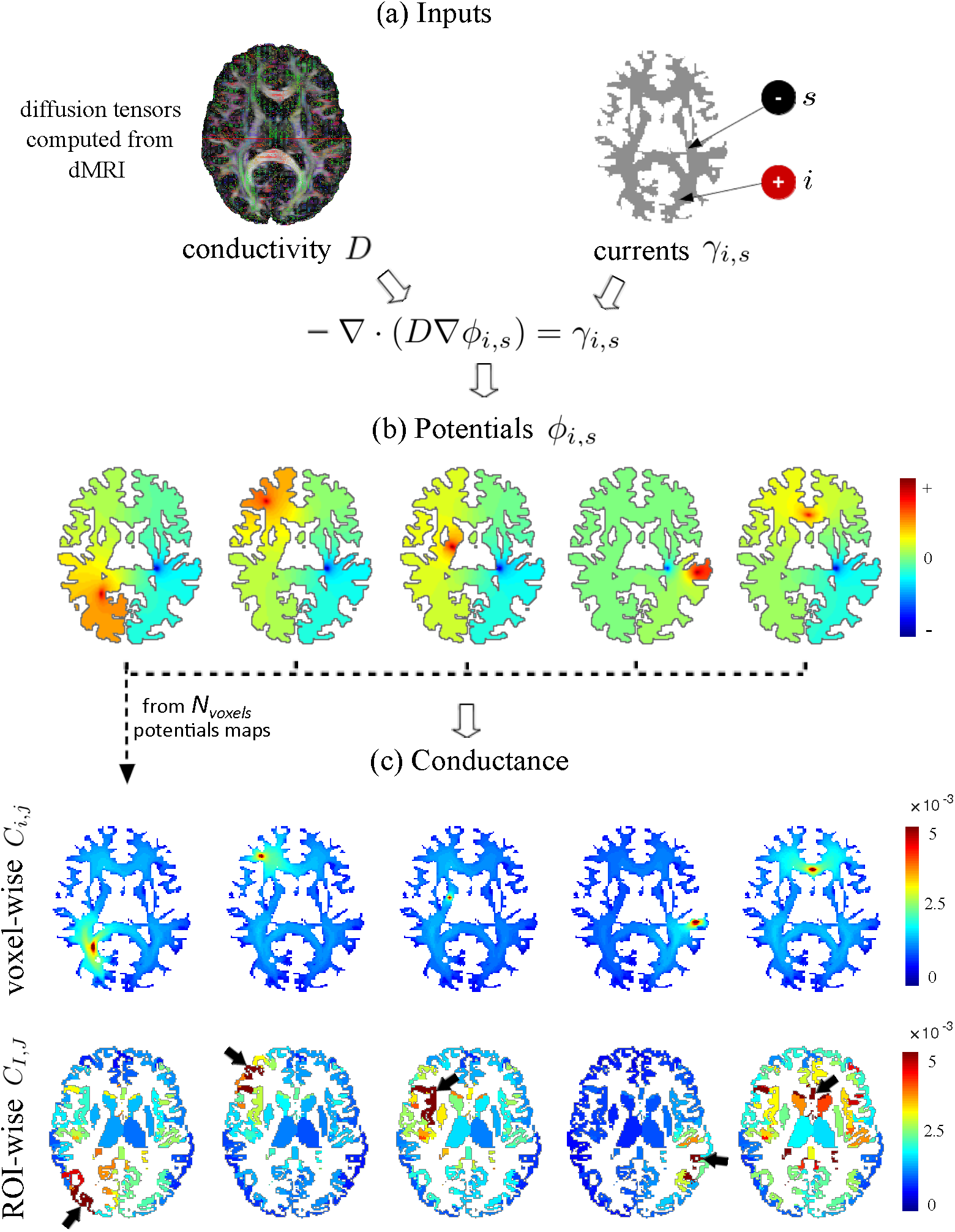
Conductance-based structural connectivity method. (a) We model brain connectivity globally using differential circuit laws, with diffusion tensors reconstructed from dMRI. (b) We first compute the potentials *ϕ*_*i,s*_, as depicted for five sources *i*(the reddest voxel) and a fixed sink *s*(the bluest voxel). (c) Next, potential maps are superposed to generate voxel-wise conductance maps *C*_*i,j*_ as shown for five voxels *i*(the reddest voxel), or ROI-wise conductance maps *C*_*I,J*_ as shown for five ROIs *I*(indicated by arrows).

We solve the PDE for a 1-ampere current (without loss of generality) between a pair of voxels *i* and *j*: *γ*_*i,j*_ = *δ*_*i*_ − *δ*_*j*_, where *δ*_*k*_(*x*) := *δ*(*x − x*_*k*_), with *x*_*k*_ the position of voxel *k* and *δ*(·) the Dirac delta. To compute ROI-wise conductance, we distribute the currents among the sets of voxels *I* and *J*(the two ROIs) as: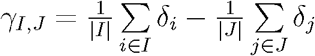

### 2.1 Efficient computation of potentials

For each source/sink current configuration *γ*_*i,j*_, we solve the PDE^1^ in Eq. (1) to find the potentials (see Fig. 1). We first discretize the linear diffusion term *-∇·*(*D∇ϕ*_*i,j*_) and *γ*_*i,j*_ (see Appendix A for details on the discretization) and write it in the matrix form *Mϕ*_*i,j*_ = *γ*_*i,j*_, which we then invert to solve *ϕ*_*i,j*_ = *M*^-1^ *γ*_*i,j*_. We use the Neumann boundary condition, ∇ *ϕ*_*i,j*_ **e** = 0 for all points on the boundary, where **e** denotes the (typically exterior) normal to the boundary.

Given *N* voxels (or ROIs), we need a potential map for each pair of voxels, leading to *N*^2^ potential maps, hence an intractable problem. However, by exploiting the superposition principle that is a consequence of the linearity of the derivative operator, we reduce the number of necessary potential maps to only *N*. We choose a reference sink voxel, *s*, and compute *N* potential maps, *ϕ*_*i,s*_, *i*= 1, *…, N*, with *i* the source and *s* the fixed sink. Next, note that for a pair (*i, j*), we can compute the potential map *ϕ*_*i,j*_ by linearly combining the potential maps between *i* and *j* and the reference sink voxel (*ϕ*_*i,s*_ and *ϕ*_*j,s*_), since:

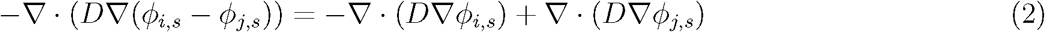

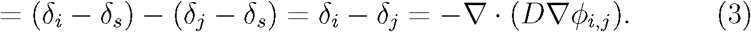

Given that we consider the same boundary conditions for all the PDEs, we now compute the potential map for any pair simply as *ϕ*_*i,j*_ = *ϕ*_*i,s*_ − *ϕ*_*j,s*_. Note that in the computation of *ϕ*_*i,j*_ for all pairs (*i, j*), the inverse matrix *M*^-1^ needs to be computed and saved in memory only once.

### 2.2 Conductance as a measure of connectivity

The conductance between two points can be computed with Ohm’s law as the ratio of the current to the potential difference. In our case, we set a 1-ampere current between two voxels (or ROIs) *i* and *j*, and the potential difference is *ϕ*_*i,j*_(*x*_*i*_) − *ϕ*_*i,j*_(*x*_*j*_). The conductance is therefore computed, voxel-wise, as:

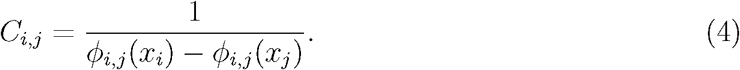

Finally, per the aforementioned superposition principle, the conductance between any pair of voxels (*i, j*) is computed as:

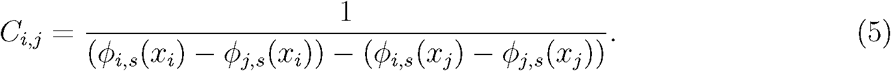

For ROI-wise connectivity, given that the potential map between the two ROIs is *ϕ*_*I,J*_ = *ϕ* _*I,s*_ *– ϕ* _*J,s*_, where 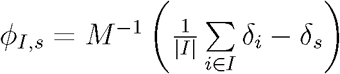 we have:

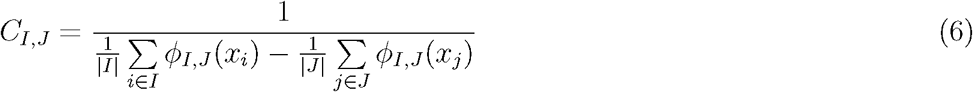

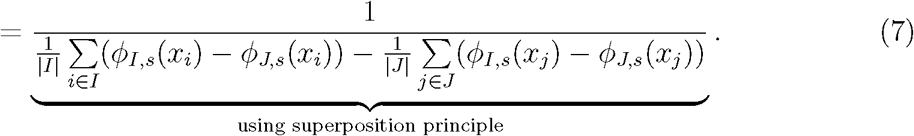

High conductance (i.e. low resistance) between two points indicates a high degree of connectivity. Note in equations (5) and (7) that, as expected, the conductance is symmetric with respect to the pair of voxels or ROIs, i.e., *C*_*i,j*_ = *C*_*j,i*_ and *C*_*I,J*_ = *C*_*J,I*_. As shown in Fig. 1(c), by setting the conductivity (D) to zero outside the white-matter mask, we can compute voxel-wise conductance maps from a single voxel (in red) to the rest of the brain, as well as ROI-wise conductance maps from a single ROI (in red, indicated with arrow) to all the other ROIs^2^. Note that the ROIs are all at least weakly connected, given that indirect connections are all considered. Nevertheless, these maps can be thresholded to keep only connections stronger than a desired amount.

### 2.3 Comparison to standard connectivity computation

In standard connectomic approaches, tractography is often performed after dMRI reconstruction to extract streamlines that represent connections between different voxels of the image (see Fig. 2, right). Connectivity matrices are then generated from the streamlines between different ROIs that are previously determined via segmentation of the brain. Various connectivity measures can be used to quantify these connections, leading to different connectivity matrices, e.g.: plain count of the streamlines, number of the streamlines normalized by the median length, number of tracts *passing* through the ROI or only those *ending* in the ROI, etc. The optimal choice of the type of connectivity matrix usually depends on the tractography algorithm used (choice of seeds, deterministic/probabilistic, whether sharp turns are allowed, etc.). In our approach, however, only the conductivity input will vary. This reduces the number of decisions to make (parameters to tune) in the computation of the connectivity matrix.

**Figure 2:**
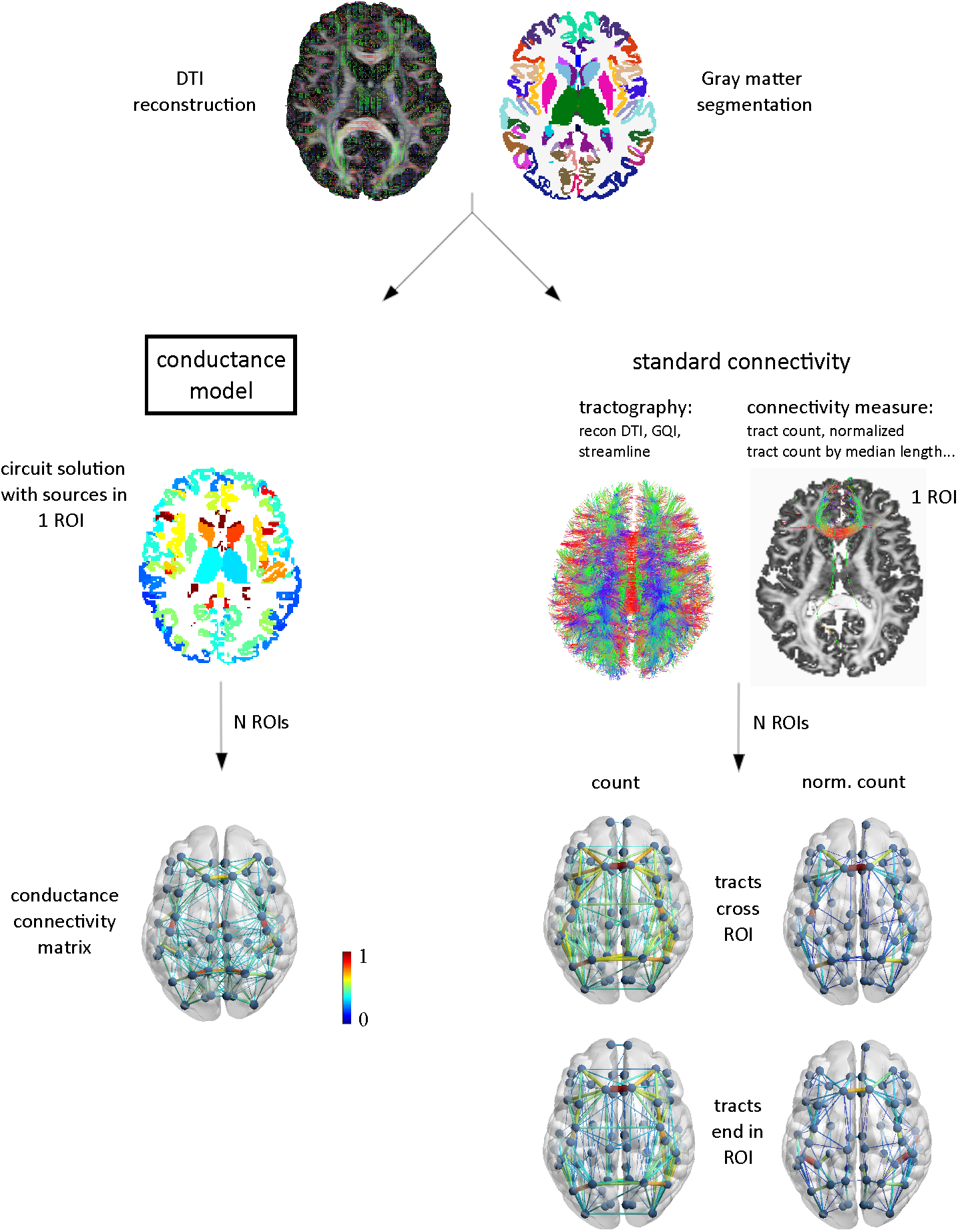
Comparison of the structural connectivity computation by the proposed method vs. standard tractography. The diffusion tensors and the white and gray matter segmentation and parcellation are given to both connectivity methods. In our conductance connectivity approach, the connectivity matrix is computed by solving a PDE for each ROI. In the standard streamline connectivity approach, tractography is performed on the tensors or higher-order distributions and connectivity matrices are computed according to certain criteria. Brain connectivity (**C**) in this figure is averaged across HCP subjects and visualized as log(1 + **C**) using the BrainNet Viewer (www.nitrc.org/projects/bnv) (Xia et al., 2013).

## 3. Data analysis

We used two publicly available datasets in our analysis, from the WashU-UMN Human Connectome Project (HCP) (Van Essen et al., 2013) and the second phase of Alzheimer’s Disease Neuroimaging Initiative (ADNI-2) (Jack et al., 2008; Beckett et al., 2015). These two datasets allowed us to evaluate the proposed method from different aspects: HCP data was used to measure the relationship between structural and functional connectivities, and ADNI-2 enabled the assessment of the utility of our measure in discriminating healthy and diseased populations. We preprocessed both datasets similarly.

### MR processing

We performed tissue segmentation and parcellation of the cortex into ROIs using FreeSurfer^3^ (Fischl, 2012). The parcellation used in this work is the Desikan-Killiany atlas (Desikan et al., 2006).

### Diffusion MRI processing

The diffusion preprocessing pipeline used for ADNI-2 involved the FSL software^4^ (Jenkinson et al., 2012) and included BET brain extraction (Smith, 2002) and EDDY (Andersson and Sotiropoulos, 2016) for eddy current and subject motion correction. For HCP, there were more steps involved (Glasser et al., 2013), namely: B0 intensity normalization, EPI distortion correction with TOPUP (Andersson et al., 2003), eddy current and subject motion correction with EDDY, and gradient non-linearity correction. From the preprocessed dMRI images, we reconstructed the diffusion tensors using the Diffusion Tensor Imaging (DTI) (Basser et al., 1994) reconstruction of DSI Studio^5^, which we then used as input to our conductance approach. To compare with standard approaches (see Fig. 2), we also ran streamline (SL) tractography (Yeh et al., 2013) using DTI, for direct comparison with our approach, and using generalized q-sampling imaging (GQI) (Yeh et al., 2010), which, as opposed to DTI, can model multiple axon populations per voxel. We generated 10000 fiber tracts and used default values for the rest of the parameters. Then, we computed connectivity matrices according to various connectivity conventions: plain tract count, tract count normalized by the median length, both considering tracts *passing* through the ROI or *ending* in the ROI.

The computational cost of the proposed conductance-based connectivity method, in terms of memory and runtime, depends on the number of voxels in the white- and gray-matter mask. The inversion of the matrix *M* is computationally expensive, thus so is our method compared to standard streamline approaches. For instance, for an ADNI-2 subject with diffusion images of dimensions 256*×* 256*×* 59 and approximately 120000 voxels in the mask, the conductance method needed up to 13GB of RAM and took approximately 6 minutes to run on a computer with 1 core (Intel(R) Xeon(R) CPU E5-2697 v3 @ 2.60GHz). For an HCP subject with diffusion images of dimensions 145 *×*174 *×*145 and approximately 570000 voxels in the mask, the conductance method needed up to 94GB of RAM and took approximately 2 hours. DSI Studio deterministic tractography, on the other hand, took approximately 5 seconds and used 4.5GB of RAM to generate 10000 streamlines from the tensors and compute the connectivity matrix, for a subject from either of the two datasets.

### rs-fMRI processing

The rs-fMRI data, already projected to the surface and preprocessed as described by Glasser et al. (2013), was detrended (only linear trends removed), bandpass-filtered at 0.01-0.08Hz, and smoothed with a kernel with a full width at half maximum of 6mm. We stacked four sessions of rs-fMRI data, and computed the matrix of Pearson’s correlations among the ROIs. Note that we used rs-fMRI only in our HCP experiments.

## 4. Results

We evaluated the performance of the proposed conductance measure through experiments that answer two different questions:

1. Relationship between structural and functional connectivity (Section 4.1): Is our structural connectivity measure, which considers both direct and indirect connections, more correlated to functional connectivity than standard measures are?
2. Discrimination of healthy and diseased populations (Section 4.2): Does our measure provide additional information that better distinguishes different stages of AD?

In the following subsections, we attempt to answer these questions using the HCP and ADNI-2 datasets, respectively.

### 4.1. HCP dataset

We evaluated our method on 100 subjects from the publicly available WashU-UMN HCP dataset (Van Essen et al., 2013), containing rs-fMRI and dMRI data, while comparing our approach with standard SL tractography in regards to the relation to functional connectivity.

#### 4.1.1 Relationship with functional connectivity

To test the hypothesis that indirect connections can explain some of the variability between structural and functional connectivity, we compared the structural connectivity matrices – computed from our conductance and the standard approaches – to the functional connectivity matrix. We computed the Pearson’s correlation coefficient between the elements of the structural and functional connectivity matrices per subject, and then compared the distribution of these correlation values, as depicted in Fig. 3(a) and reported in Table 1. As can be seen, functional connectivity is much more strongly correlated with the proposed conductance-based structural connectivity than with all the conventional SL-based connectivity methods. Next, to make sure that our method makes use of the actual white-matter fiber orientations, and not just the physical distance between ROIs, we repeated the experiments using isotropic tensors; once with a constant isotropic tensor magnitude (denoted by ‘distance’), and again by weighting the isotropic tensor with mean diffusivity (denoted by ‘MD-w distance’). Figure 3(b) shows the histogram of the correlations of functional connectivity with two SL-based connectivity approaches (DTI, GQI) subtracted (subject-wise) from correlations of functional connectivity with the proposed conductance-based connectivity. This figure, as opposed to Figure 3(a), is a *paired* comparison and preserves the subject-wise information by subtracting the connectivity values subject by subject, and then illustrating the histogram of the differences. The positive range of values shows how much more the conductance connectivity is correlated with the functional connectivity than the conventional SL-based metrics are. A two-tailed paired *t*-test between these two distributions revealed a statistic of *t*= 37 and a significance value of *p*= 10^−60^ in the DTI case, and *t*= 35 and *p*= 10^−58^ in the GQI case.

**Table 1:** Cross-subject mean correlation of functional and the proposed conductance-based structural con<u>nectivities with all structural connectivity metrics. See the legend of Fig. 3</u> for keyword definitions.

|  | conductance | DTI SL |  |  |  | GQI SL |  |  |  |
| --- | --- | --- | --- | --- | --- | --- | --- | --- | --- |
|  |  | end |  | pass |  | end |  | pass |  |
|  |  | count | ncount | count | ncount | count | ncount | count | ncount |
| functional | **0.427** | 0.163 | 0.152 | 0.165 | 0.192 | 0.141 | 0.138 | 0.175 | **0.195** |
| conductance | 1 | 0.304 | 0.346 | 0.353 | **0.439** | 0.251 | 0.336 | 0.356 | **0.453** |

**Figure 3:**
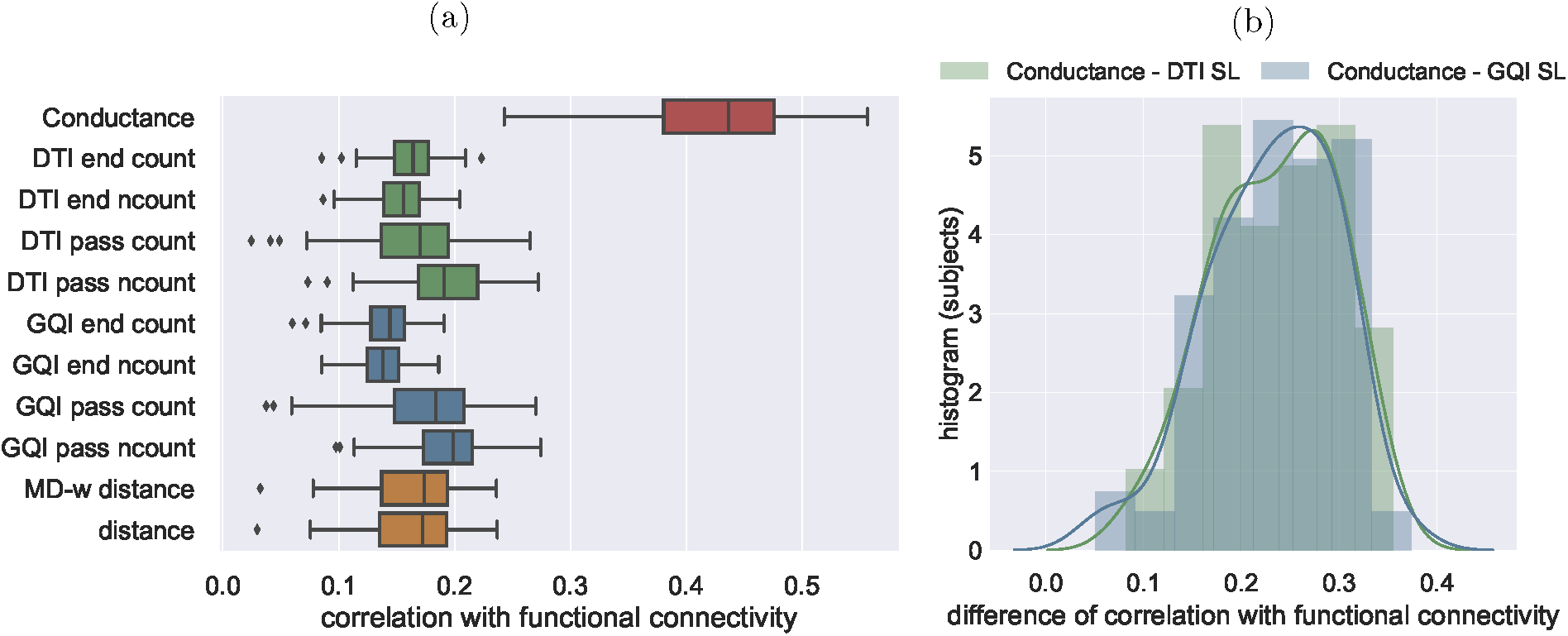
Correlation of structural and functional connectivities considering different approaches and metrics. Distribution of correlations across 100 subjects. SL-based connectivity measures computed using DTI and GQI are compared: ‘count’ means plain count, ‘ncount’ means normalized fiber count by the median length, ‘end’ means that only fibers *ending* in the ROI are considered, and ‘pass’ that fibers *passing* through an ROI are also considered. The ‘distance’ method is our approach ran with constant isotropic tensors, and the ‘MD-w distance’ is our approach ran with isotropic tensors weighted by the mean diffusivity. For each box, the central line is the median, the left/right edges are the 25^th^ and the 75^th^ percentiles, the whiskers extend to the most extreme data points not considered outliers, and the outliers are plotted individually. Histogram of the difference of structural/functional correlation, for conductance minus DTI SL and GQI SL tractographies. The SL-based connectivity measure giving the highest correlation with functional connectivity is used: tract count normalized by median length, considering tracts *passing* through an ROI.

#### 4.1.2. Modeling of inter-hemispheric connections

One of the well-known issues in standard tractography methods is the mismodeling of inter-hemispheric connections, as such long tracts can be mistakenly cut short (Sinke et al., 2018). As can be seen in Table 2, conventional structural connectivity methods model a low quantity of cortical inter-hemispheric (only 6% of all) connections. Our conductance method results in a higher ratio of inter-hemispheric connections, close to that of functional connectivity measures and the MD-w distance.

**Table 2:** HCP inter-hemispheric connection ratios with respect to all connections, for different connectivities. This ratio was computed as the sum of ROI-to-ROI connectivity values across inter-hemispheric connections, divided by the sum of all ROI-to-ROI connectivity values. Reported results were averaged across 100 subjects.

| connectivity | ratio of cortical inter-hemispheric to all |
| --- | --- |
| MD-w distance connections | $0.3278 \pm 0.0131$ |
| functional correlations | $0.3356 \pm 0.0310$ |
| conductance model connections | $0.2849 \pm 0.0071$ |
| GQI SL pass ncount connections | $0.0613 \pm 0.0119$ |

Next, we again considered the correlation of the different structural connectivity measures with functional connectivity, but this time separately for three connection groups of cortical inter-hemispheric, cortical intra-hemispheric, and all subcortical connections, with the results provided in Table 3 and illustrated in Fig. 4. Regarding the cortical inter-hemispheric connections, we observe a much lower correlation of the SL-based measures compared to the conductance-based measure, with no overlap. Note that our measures of distance have low correlations as well. This difference decreases for intra-hemispheric connections, although conductance-based connectivity still correlates much more than the rest. Although extreme values in the results by our method overlap with those by some DSI Studio measures, their 25^th^/75^th^ percentiles are still far from each other. As for subcortical connections, we observe that the distance correlates much more with functional connectivity than SL-based connectivity measures do, and that the conductance-based connectivity has a very large standard deviation. As a result, we can conclude that the deviation for our method in Fig. 3 originates mostly from subcortical connections.

**Table 3:** Results of two-tailed paired *t*-tests between the distributions of the structure-function correlations computed by the conductance connectivity and the streamline deterministic tractography using normalized count of tracts *crossing* an ROI.

|  | all connections | cortical<br>inter-hemispheric | cortical<br>intra-hemispheric | subcortical |
| --- | --- | --- | --- | --- |
| DTI | $t = 37, p = 10^{-60}$ | $t = 43, p = 10^{-65}$ | $t = 34, p = 10^{-56}$ | $t = 12, p = 10^{-21}$ |
| GQI | $t = 35, p = 10^{-58}$ | $t = 41, p = 10^{-64}$ | $t = 33, p = 10^{-55}$ | $t = 15, p = 10^{-26}$ |

**Figure 4:**
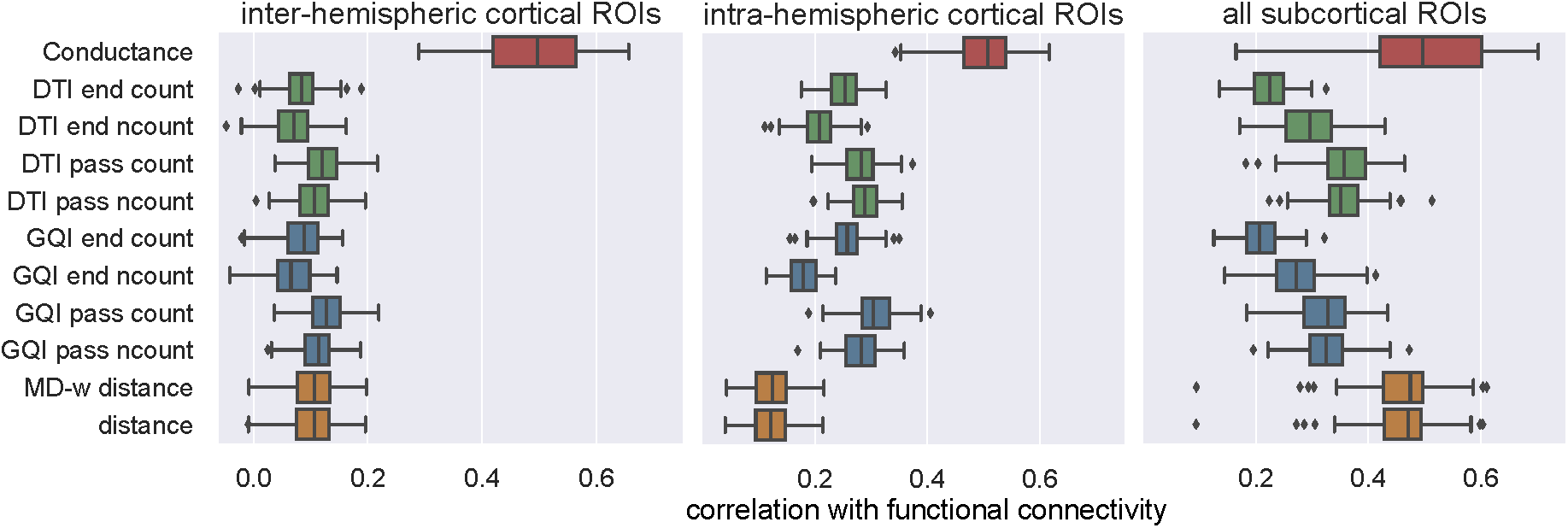
Structure/function correlation for three groups of connections: cortical inter-hemispheric, cortical intra-hemispheric and all subcortical connections. Comparison of conductance model, the SL-based connectivity measures and *distance* and *MD-w distance*. For each box, the central line is the median, the left/right edges are the 25^th^ and 75^th^ percentiles, the whiskers extend to the most extreme data points not considered outliers, and the outliers are plotted individually.

**Figure 5:**
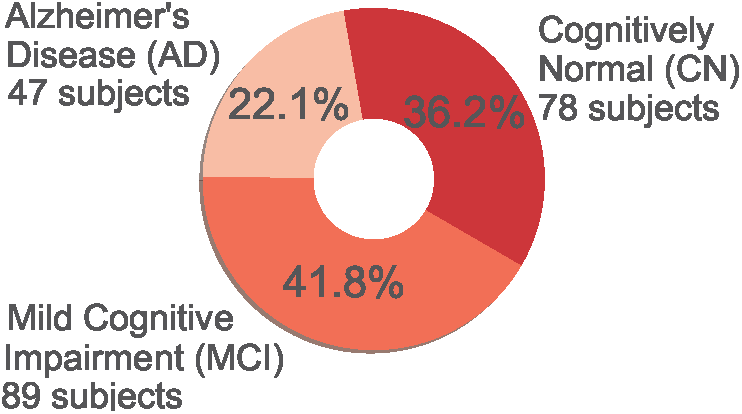
ADNI-2 data demographics.

### 4.2. ADNI-2 dataset

Next, we tested the power of our global conductance-based structural connectivity measure in the detection of white-matter differences in populations with different stages of AD and in the classification of dMRI images according to the AD stage. A more accurate modeling of structural connections could lead to a better understanding of the progression of this and other diseases, and to the discovery of new imaging biomarkers for disease study.

We analyzed 213 subjects of the ADNI-2 dataset (http://adni.loni.usc.edu) in different stages: 78 cognitively normal (CN), 89 with mild cognitive impairment (MCI), and 47 with AD dementia (Fig.5) The ADNI was launched in 2003 as a public-private partnership, led by Principal Investigator Michael W. Weiner, MD. The primary goal of ADNI has been to test whether serial MRI, positron emission tomography, other biological markers, and clinical and neuropsychological assessment can be combined to measure the progression of MCI and early AD.

#### 4.2.1. Disease stage classification

We assessed the suitability of our conductance-based connectivity matrix for disease stage classification, which would allow inferring the disease stage from connectivity measures derived from dMRI. We performed the classification of the elements of the ROI-to-ROI connectivity matrix with a Random Forest classifier with balanced class weight (Pedregosa et al., 2011). We cross-validated with a shuffle split cross-validation strategy with 200 splits considering a 10% of the data for testing (Varoquaux et al., 2017).

We classified each pair of stages, and then all stages together, and report the results in Fig. 6. We compare our conductance measure to different conventional SL-based connectivity measures, using DTI and GQI reconstruction methods. Note that, in general, classification performance is affected by class size, even though we used balanced weights in the classification. In all cases, our conductance measure has a higher prediction accuracy than conventional structural connectivity measures do, especially in the case of CN vs AD. When classifying all stages simultaneously, one must keep in mind that since we have 3 classes, a prediction accuracy of 0.5 is still informative. For the case of all-stage classification, Fig. 7 shows the confusion matrices for our conductance measure and the conventional structural connectivity measure (here only considering the measure from GQI with SL tractography, counting the number of tracts passing through an ROI normalizing by the median length). We observe that our conductance method correctly classifies 45% of the AD cases whereas the conventional SL connectivity measure does only 12%. For the rest of the disease stages, the two methods had comparable performance.

**Figure 6:**
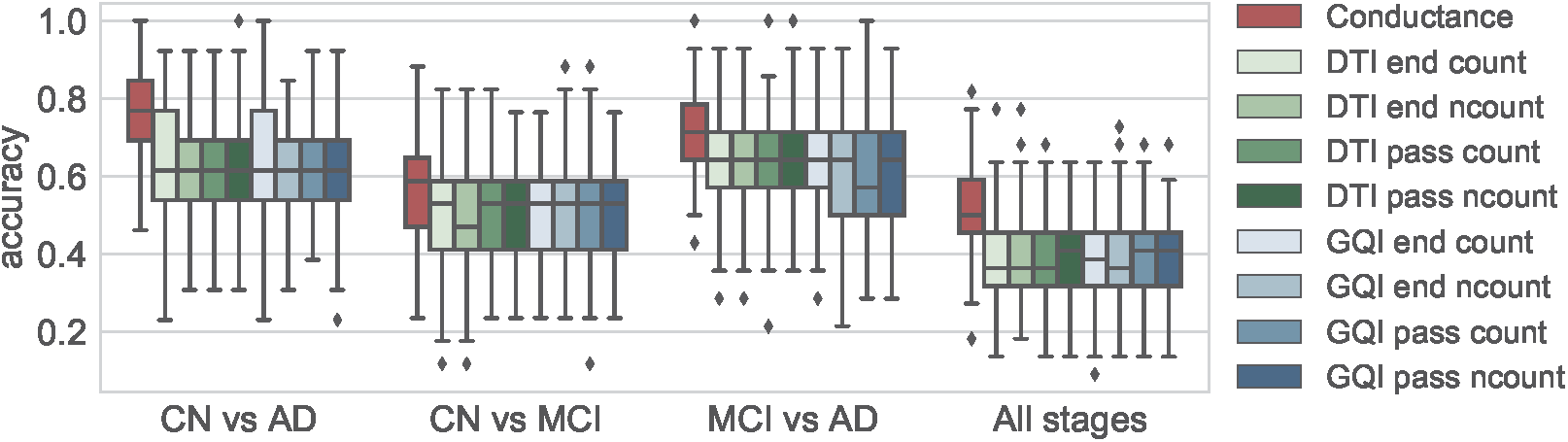
Prediction accuracies of differenctomstpages of AD, when using a Random Forest classifier, with balanced weights. The cross-validation used was: Shuffle Split with 200 splits considering 10% of the data for testing. See the legend of Fig. 3 for keyword definitions.

**Figure 7:**
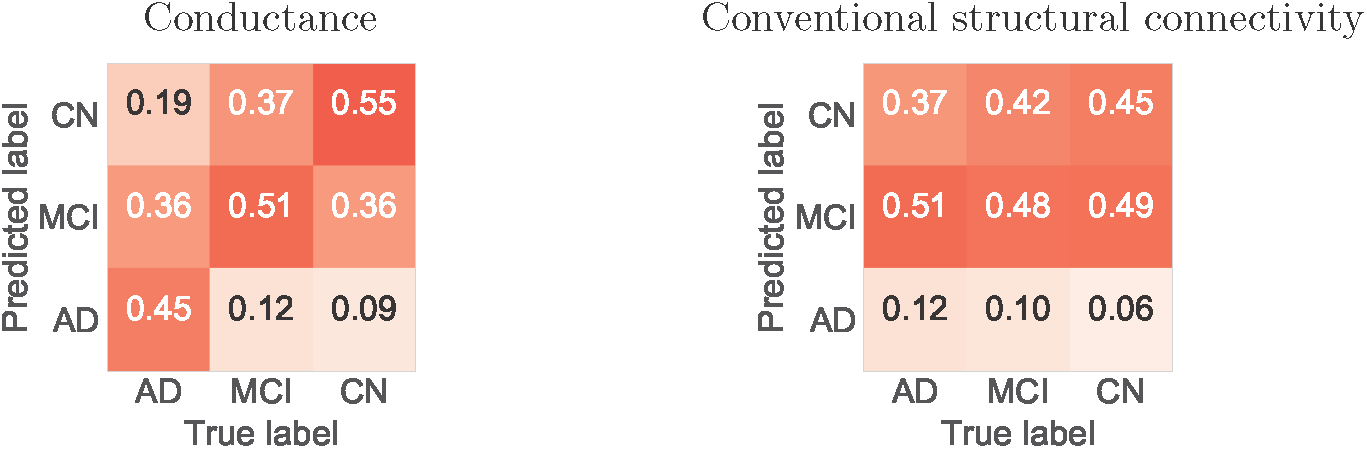
Confusion matrices and prediction accuracy in the classification of disease stages using a Random Forest classifier on the conductance method connectivity matrix and the conventional SL connectivity matrix with GQI, and the metric fiber count normalized by the median tract length.

#### 4.2.2. Pairwise group comparison

Next, we searched for pairs of regions where the structural connectivity significantly differed between the CN and AD groups. Figure 8 shows, for each pair of ROIs, the *p*- values resulting from a 2-sample *t*-test between the AD and CN groups. Connections with significant differences (*p <*0.05, after Bonferroni correction) between the AD and CN groups for the conductance measure (Fig. 8(a)) involve regions that are known to be affected by AD, namely hippocampus and amygdala (Barnes et al., 2006; Prestia et al., 2011). Regarding differences between the MCI and AD groups, only the connections of right pars opercularis with insula and amygdala were significant, and when considering the CN and MCI groups, no connection was significantly different.

**Figure 8:**
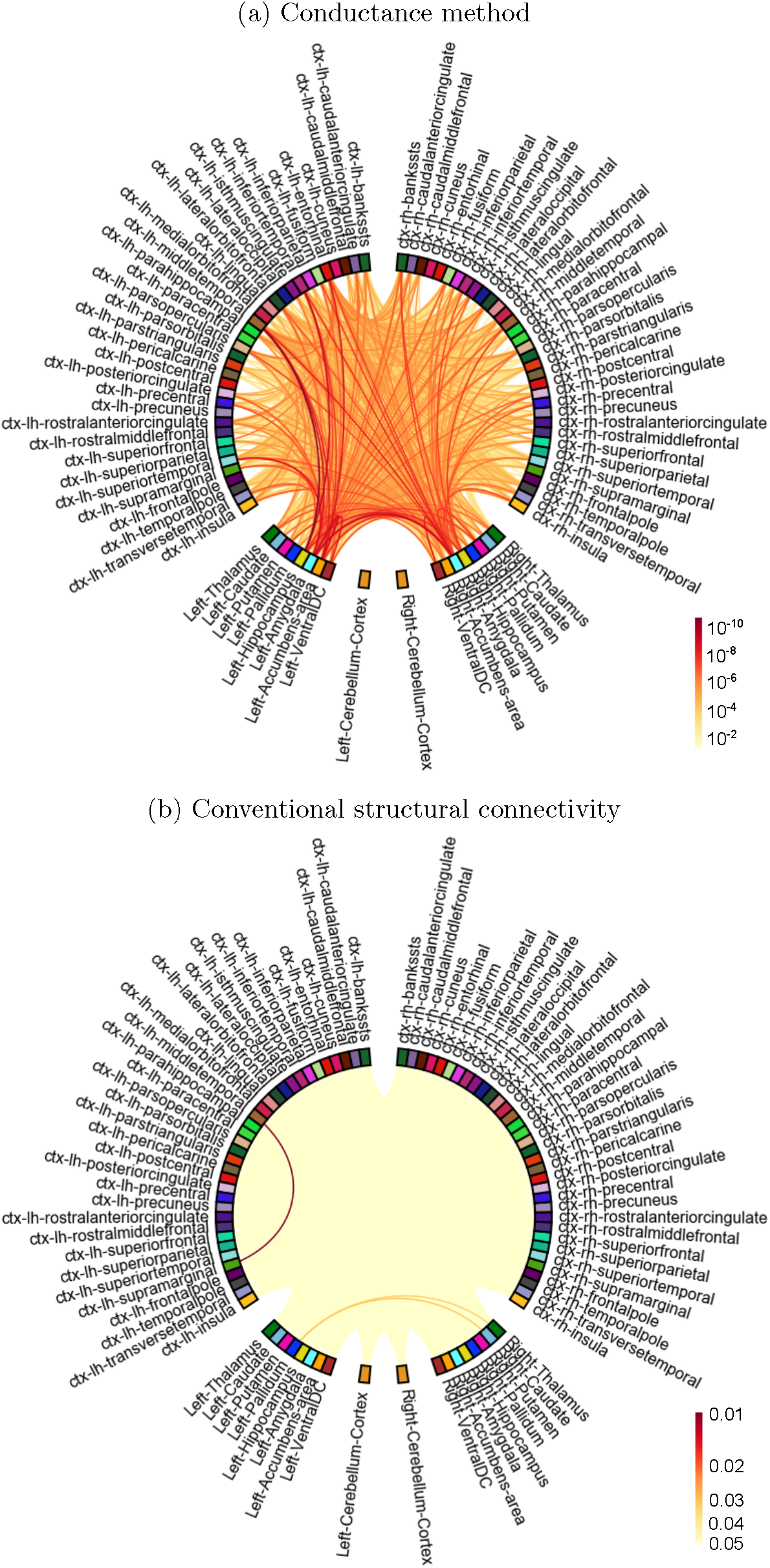
*p*-values showing significant differences in connectivity per ROI pair between the AD and CN groups, when using (a) the proposed conductance method and (b) a conventional structural connectivity measure (DTI measure counting the number of fibers passing through an ROI).

We performed the same analysis on the conventional SL connectivity measures (Fig. 8(b)), and only some of the DTI measures revealed a few significantly different connections in the comparison of AD and CN groups. DTI counting the number of fibers *passing* through an ROI and the same normalized by the median length revealed differences in a few connections, as depicted in Fig. 8(b). When counting the number of fibers either *passing* through or *ending* in an ROI, only the left middle-temporal to supramarginal connection was significantly different. No differences were found in the other pairs of groups, and none with any of the GQI SL measures.

Prior to Bonferroni correction, both methods suggested many significant connections between all pairs of stages. However, only the proposed method revealed significant *p*-values in all cases after Bonferroni correction. This suggests that our method provides a higher statistical power and might potentially reveal a statistically significant effect with a smaller sample size.

## 5. Discussion

We have proposed a new approach to measure structural brain connectivity. Directly from the reconstructed dMRI data, our approach accounts for indirect brain connections that would not otherwise be inherently considered by standard SL techniques. The proposed methodology is solved globally, and its primary advantage is that it considers all possible pathways when quantifying connectivity. As such, however, it may also be including some implausible paths, as would also be the case for the conventional SL tractography-based connectivity measures. Nonetheless, we expect the greater length of erroneous pathways, such as the ones that involve jumping from one true path onto another, to reduce the impact of such paths. It is important to note that, although the diffusion tensor has been shown to be related to the physical conductivity tensor (Tuch et al., 2001), we do not suggest either that the diffusion tensors are precise estimates of the conductivity, or that the physical electrical conductance is an appropriate model for the brain’s biological wiring. We only seek to take advantage of the mathematical framework provided by electromagnetic models to conveniently account for indirect white-matter connections, without implying that such a connectivity measure would explain any physical properties of the brain.

Another advantage of our method is that we have fewer decisions to make and parameters to tune in the connectivity analysis pipeline. Standard tractography-based connectivity analysis includes different steps that require a certain decision making when quantifying connectivity, such as: modeling of one or several fiber bundle populations, the choice of probabilistic or deterministic tractography, number of seeds, number of streamlines, the allowed turning angle, how we count the tracts, etc. This introduces a large amount of variability in the results, in a field where the scarcity of ground truth is one of the biggest challenges (Cheng et al., 2012; Maier-Hein et al., 2017). Our methodology, however, computes connectivity from the diffusion tensors without the need for any parameter tuning. It is worth noting that, even though there is some variability in the results by conventional connectivity methods, our method consistently outperformed them.

We have shown in Section 4.1 that by using the proposed conductance method, which accounts for indirect pathways, one can compute structural connectivity measures that are significantly more correlated with functional connectivity than by using more standard approaches, with the latter showing a very low correlation in line with existing literature (Fjell et al., 2017). This supports the hypothesis on the role of indirect connections in the relationship between functional and structural connectivity; part of the variance between these two connectivity measures lies within indirect connections, usually not accounted for in structural connectivity (Honey et al., 2009; Deligianni et al., 2011). Although the conductance measure was significantly more correlated with functional connectivity than standard SL measures were, the mean correlation was merely 0.43 (see Fig. 3). The reason for this is that structural and functional connectivities describe different processes: while the former represents physical connections, the latter reflects how synchronously different regions in the brain function and considers such synchrony as a surrogate for connectivity. Nevertheless, increased similarity to functional connectivity – when considering all connections or only a subgroup of them (see Fig. 4) – implies that the proposed measure may be more informative about the brain function.

We observed through experiments that our approach reveals more cortical inter-hemispheric connections (see Section 4.1.2). These connections, which are mostly – but not always (Roland et al., 2017) – through the corpus callosum (Zarei et al., 2006), have proved challenging for standard tractography methods (e.g., due to sensitivity to head motion) (Yendiki et al., 2014). In fact, inter-hemispheric connections have been shown to be affected by disease (e.g., AD) and correlated with clinical changes in some disorders (Wang et al., 2013, 2015; Saar-Ashkenazy et al., 2016; Xue et al., 2018). Further investigation with experiments on datasets with known ground truth, e.g. from chemical tracing (Grisot et al., 2018), would be required to find out whether better revealing of inter-hemispheric connections by our method compared to deterministic streamline tractography is only a consequence of modeling indirect connections, or also of better capturing of some hard-to-detect direct ones.

The proposed conductance measure has also proven useful (see Section 4.2) in distinguishing normal and AD subjects and in better classification among AD stages. This could potentially lead to dMRI-derived biomarkers for early detection of AD.

Our approach could be employed in a more general-purpose global connectivity analysis, as an alternative to classic tractography. However, the conductance values may not fit the definition of either streamlines or probability measures. As we have shown in Section 2, the proposed method can also enable the computation of voxel-wise and thus parcellation-independent connectivity (Moyer et al., 2017), which allows us to compute the connectivity between any pair of points in the white matter.

One of the limitations of the proposed conductance measure is that it derives from tensors, and therefore does not make use of the information provided by higher-order models. Even so, we make use of the whole tensor (not just the main direction of the axon population), which is informative in distinguishing highly anisotropic fibers from the cases where fiber crossing or fanning is present. Therefore, our tensors are actually used as low-order orientation distribution functions (ODFs). Importantly, we have shown here that our approach outperforms the conventional SL-based method, even when the latter uses high-order ODFs. In addition, our approach can be extended to exploit multiple axon population models to deal with crossing and kissing fibers, using a multi-tensor approach or general ODFs (Seunarine and Alexander, 2014; Aganj et al., 2015) instead. We propose a framework to implement this with higher-order models in Appendix B.

## 6. Conclusion

In this work, we have proposed a new structural connectivity measure that is directly derived from the dMRI data and accounts for all possible pathways when quantifying connectivity between a pair of regions. The method is global, does not produce locally optimal solutions, and has no parameters to be tuned by the user. Using this methodology, we computed structural connectivity measures that were significantly more correlated with functional connectivity, when compared to more standard approaches. In the study of diseased populations, our novel connectivity measure better classified different stages of Alzheimer’s disease. Our results suggest that the proposed method provides new information that is not accounted for in standard streamline-based connectivity measures, and highlights its potential for further development. Future work includes additional validation, e.g. with chemical tracing as the ground truth, which may clarify the nature of the new information provided by our method.

## Acknowledgments

This research was supported by the BrightFocus Foundation (A2016172S). Additional support was provided by the National Institutes of Health (NIH), specifically the National Institute of Diabetes and Digestive and Kidney Diseases (K01DK101631, R21DK108277), the National Institute for Biomedical Imaging and Bioengineering (P41EB015896, R01EB006758, R21EB018907, R01EB019956, R01EB021265), the National Institute on Aging (AG022381, 5R01AG008122-22, R01AG016495-11, R01AG016495), the National Center for Alternative Medicine (RC1AT005728-01), the National Institute for Neurological Disorders and Stroke (R01NS052585, R21NS072652, R01NS070963, R01NS083534, U01NS086625), the National Institute of Mental Health (U01MH108168), and the NIH Blueprint for Neuroscience Research (U01MH093765), part of the multi-institutional Human Connectome Project. Computational resources were provided through NIH Shared Instrumentation Grants (S10RR023401, S10RR019307, S10RR023043, S10RR028832).

HCP data were provided [in part] by the Human Connectome Project, WU-Minn Consortium (Principal Investigators: David Van Essen and Kamil Ugurbil; 1U54MH091657) funded by the 16 NIH Institutes and Centers that support the NIH Blueprint for Neuroscience Research; and by the McDonnell Center for Systems Neuroscience at Washington University.

ADNI data collection and sharing was funded by the Alzheimer’s Disease Neuroimaging Initiative (ADNI) (National Institutes of Health Grant U01 AG024904) and DOD ADNI (Department of Defense award number W81XWH-12-2-0012). ADNI is funded by the National Institute on Aging, the National Institute of Biomedical Imaging and Bioengineering, and through generous contributions from the following: AbbVie, Alzheimer’s Association; Alzheimer’s Drug Discovery Foundation; Araclon Biotech; BioClinica, Inc.; Biogen; BristolMyers Squibb Company; CereSpir, Inc.; Cogstate; Eisai Inc.; Elan Pharmaceuticals, Inc.; Eli Lilly and Company; EuroImmun; F. Hoffmann-La Roche Ltd and its affiliated company Genentech, Inc.; Fujirebio; GE Healthcare; IXICO Ltd.; Janssen Alzheimer Immunotherapy Research & Development, LLC.; Johnson & Johnson Pharmaceutical Research & Development LLC.; Lumosity; Lundbeck; Merck & Co., Inc.; Meso Scale Diagnostics, LLC.; NeuroRx Research; Neurotrack Technologies; Novartis Pharmaceuticals Corporation; Pfizer Inc.; Piramal Imaging; Servier; Takeda Pharmaceutical Company; and Transition Therapeutics. The Canadian Institutes of Health Research is providing funds to support ADNI clinical sites in Canada. Private sector contributions are facilitated by the Foundation for the National Institutes of Health (www.fnih.org). The grantee organization is the Northern California Institute for Research and Education, and the study is coordinated by the Alzheimer’s Therapeutic Research Institute at the University of Southern California. ADNI data are disseminated by the Laboratory for Neuro Imaging at the University of Southern California.

BF has a financial interest in CorticoMetrics, a company whose medical pursuits focus on brain imaging and measurement technologies. BF’s interests were reviewed and are managed by Massachusetts General Hospital and Partners HealthCare in accordance with their conflict of interest policies.

## Appendix A. Discretization of the diffusion term

For the discretization of *∇* (*D∇ ϕ*) of Eq. (1), we find the values of *D* on the mid points of the mesh before computing the divergence, as such mid points are where the gradient of the potentials are computed.

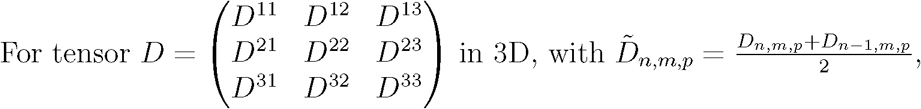

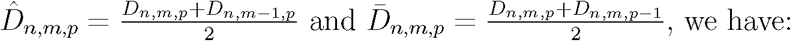

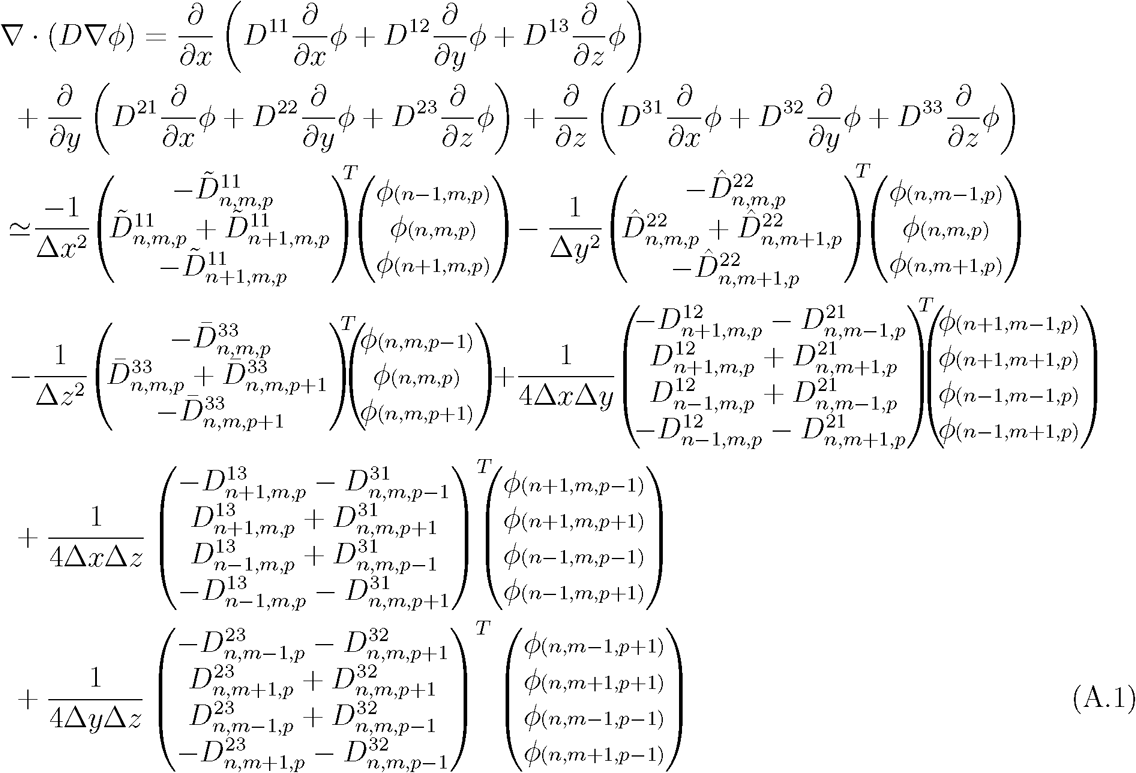

## Appendix B. Extension for high-angular-resolution diffusion imaging (HARDI)

Assuming that 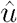 is the normalized *b*-vector and *b*= *τq*^2^ the *b*-value, with *q* and *τ* the *q*-vector and the diffusion time, the measured diffusion signal values according to the monoexponential model are

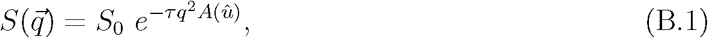

where 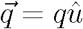 and *A* is the apparent diffusion coefficient. Note that

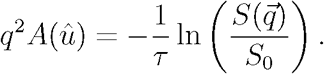

In DTI, we approximate *A* as a quadratic function:

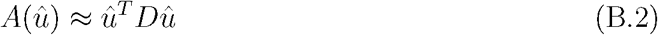

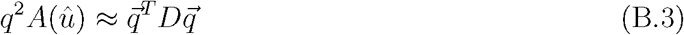

Deriving with respect to 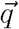 results in:

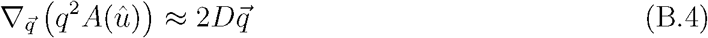

For HARDI, we can approximate the left-hand-side of Eq. (1) as follows:

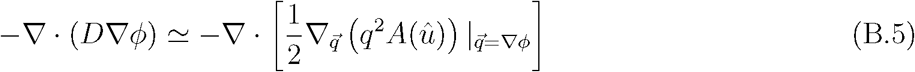

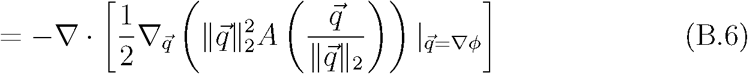

Equation (B.6) is no longer linear with respect to the potentials, and therefore Eq. (1) cannot be solved as easily with this model as in the case of DTI. Moreover, due to nonlinearity, the superposition property no longer holds either. Iterative optimization approaches may be used to approximate the solution to the PDE with Eq. (B.6).

## Footnotes

1 We used the Finite Volume MATLAB toolbox (Eftekhari and Schller, 2015), and extended it so it accepts tensor-valued D.

2 Our codes are publicly available at: www.nitrc.org/projects/conductance

3 FreeSurfer, https://surfer.nmr.mgh.harvard.edu/

4 FSL, https://fsl.fmrib.ox.ac.uk/fsl/fslwiki/

5 DSI Studio, http://dsi-studio.labsolver.org

